# Preclinical trial of avocado pulp supplementation in an L-NAME model of cardiovascular injury

**DOI:** 10.1101/2025.10.10.681762

**Authors:** Joy A.C. Amadi, Oluchi A. Amadi, Chukwuma H. Chukwu, Peter U. Amadi

## Abstract

**Background:** Endothelial dysfunction, dyslipidemia, and myocardial injury are major contributors to cardiovascular disease. Avocado (Persea americana), rich in monounsaturated fatty acids and phytochemicals, has shown lipid-lowering and anti-inflammatory properties, but its integrated effects on vascular injury remain unclear.

**Methods:** Male rats were randomized into six groups (n = 4 per group): control, avocado, L-NAME, L-NAME+drugs (metoprolol+losartan), L-NAME+avocado, and L-NAME+drugs+avocado. Morphometric indices, lipid profiles, cardiac injury enzymes, and vascular biomarkers were measured after treatment. One-way ANOVA with Tukey test assessed group differences, while contour plots and correlation networks visualized biomarker interactions.

**Results:** L-NAME treatment induced a pathological phenotype characterized by reduced feed efficiency (-40%), weight gain (-80%), and BMI (-18%), together with dyslipidemia (LDL +120%, TG +55%, TC +42%, HDL -28%), myocardial stress (troponin +70%, CK +50%, LDH +35%), and vascular activation (endothelin +350%, VCAM-1 +55%, AngII +80%; all p < 0.01). Avocado supplementation mitigated these effects: BMI and feed efficiency returned to near-control levels, LDL, TG, and TC fell by 30-45%, and troponin, CK, and LDH decreased by ∼25-30%. Endothelin, VCAM-1, and AngII were reduced by 40-55% relative to L-NAME. Network analysis revealed dense pathological correlations under L-NAME (density 0.42), simplified under avocado (0.17), and most normalized with avocado+drugs (0.09), indicating restoration of physiological biomarker independence.

**Conclusion:** Avocado supplementation attenuates L-NAME-induced vascular injury by improving metabolic efficiency, correcting dyslipidemia, reducing cardiac injury, and dampening endothelial activation, while reprogramming pathological biomarker networks toward control-like organization

**Highlights:**

- Avocado supplementation improves lipid balance, cardiac integrity, and vascular function in L-NAME–induced injury.
- Contour and network analyses reveal avocado disrupts maladaptive biomarker couplings and restores control-like organization.
- Preclinical evidence supports avocado as a nutraceutical adjunct for integrated cardiometabolic protection.

**GRAPHICAL ABSTRACT:** 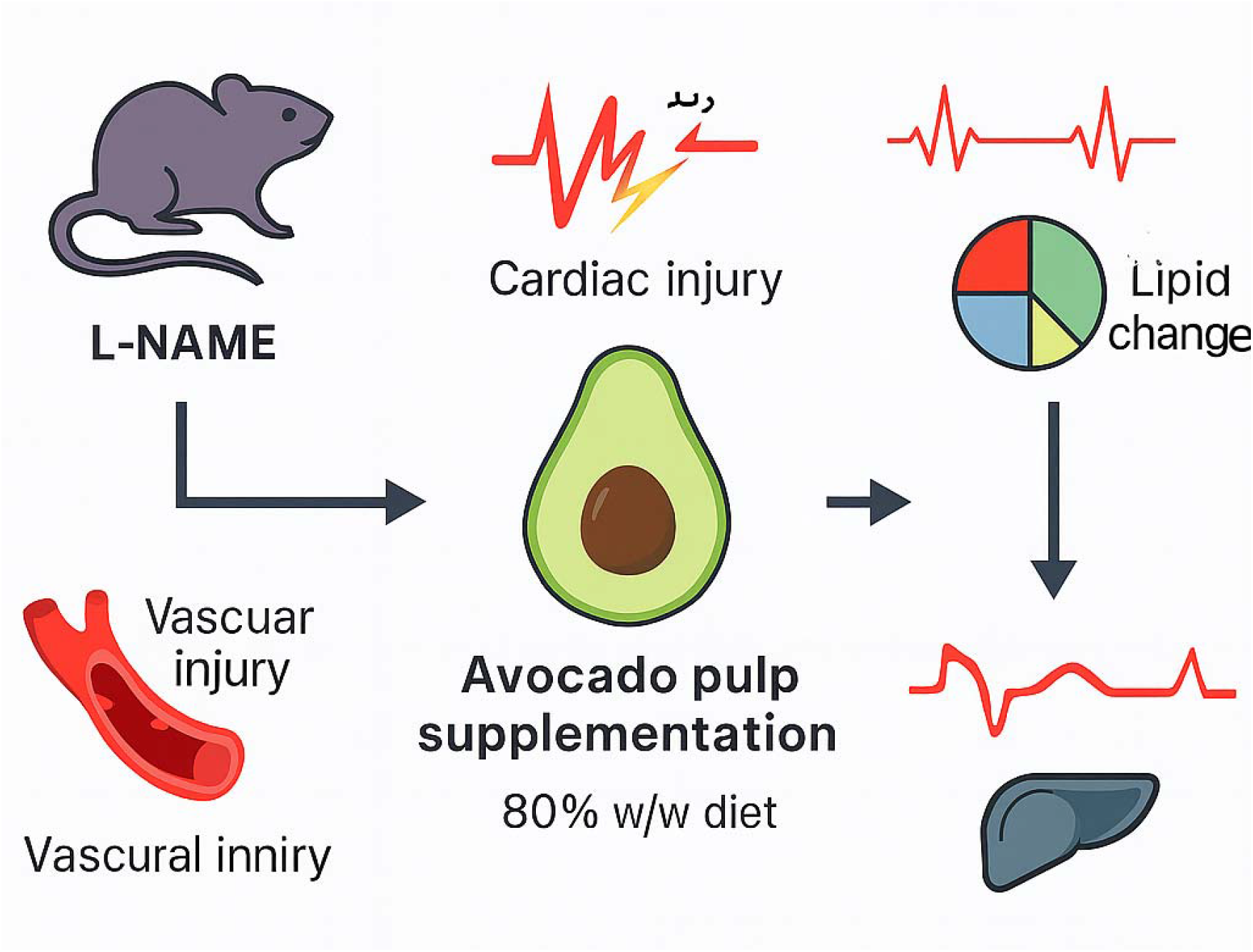

## Introduction

Cardiovascular diseases (CVDs) remain the leading cause of morbidity and mortality worldwide, accounting for an estimated 17.9 million deaths annually.^1^ Despite advances in pharmacological therapy, including antihypertensives, lipid-lowering drugs, and antiplatelet agents, the global burden of vascular dysfunction, myocardial injury, and metabolic syndrome continues to rise. Endothelial dysfunction and vascular inflammation are central drivers of this burden, promoting hypertension, atherosclerosis, and organ damage through maladaptive interactions between lipid metabolism, neurohormonal signaling, and cardiac integrity. Identifying interventions that not only target isolated risk factors but also reprogram the interconnected biomarker networks that underpin cardiovascular pathology is therefore an urgent priority.^2^ Experimental models provide insight into the integrated pathophysiology of vascular injury.

Chronic administration of Nω-nitro-L-arginine methyl ester (L-NAME), a nitric oxide synthase inhibitor, induces hypertension, endothelial dysfunction, and cardiac stress, recapitulating key features of human vascular disease.^3^ L-NAME–treated animals exhibit dyslipidemia, increased circulating endothelin, and activation of the renin–angiotensin system, along with elevated cardiac enzymes and troponin, all of which reflect vascular inflammation and myocardial injury.^4^ Importantly, these disturbances do not occur in isolation; rather, they form dense, pathological correlation networks in which lipid abnormalities reinforce vascular activation and cardiac stress.^5^ This systems-level integration highlights the limitations of single-target therapies and underscores the need for multifaceted interventions capable of reshaping pathological biomarker interactions.^6^

Dietary strategies represent a promising avenue for addressing this gap. Epidemiological and interventional studies increasingly support the role of plant-derived foods rich in unsaturated fatty acids, phytosterols, and antioxidants in lowering cardiovascular risk.^7^ Avocado (Persea americana) has emerged as a functional food with demonstrated benefits on lipid profiles, glycemic control, and vascular function.^8^ Avocado is uniquely rich in monounsaturated fatty acids, fiber, phytochemicals, and bioactive compounds with anti-inflammatory and antioxidant properties.^8–11^ Clinical studies have shown that avocado consumption can reduce LDL cholesterol, elevate HDL cholesterol, and improve endothelial reactivity.^12–14^ However, the extent to which avocado supplementation can counteract integrated vascular, cardiac, and metabolic injury in experimental models of endothelial dysfunction has not been comprehensively examined.

In this study, we used the L-NAME model of vascular injury to test whether avocado supplementation alone or in combination with standard pharmacological therapy could ameliorate cardiometabolic dysfunction. We evaluated morphometric indices, lipid profiles, cardiac enzymes, vascular biomarkers, and integrative outcomes using contour density plots and correlation network analyses. By complementing traditional statistical comparisons with systems-level visualization, we sought to determine not only whether avocado improves individual markers but also whether it remodels the pathological biomarker interactions that characterize vascular injury. We report that avocado supplementation preserved metabolic efficiency, normalized dyslipidemia, reduced cardiac injury markers, and attenuated endothelial activation. Beyond these effects, avocado disrupted maladaptive biomarker couplings, simplified pathological correlation networks, and restored control-like systems-level organization. These findings position avocado as a dietary intervention with broad, integrative cardioprotective potential, supporting its further evaluation as a nutraceutical adjunct in cardiovascular prevention and therapy.

## 2.0: Methodology

### Experimental Design and Animal Housing

Male Wistar rats (8–10 weeks old; 180–200 g) were procured from the Animal Facility of Imo State University and acclimatized for 7 days under standard laboratory conditions (22 ± 2 °C, 12 h light/dark cycle, 50–60% humidity). Rats had free access to commercial chow (UAC Nigeria Grand Cereals, Jos, Nigeria) and water ad libitum. All procedures complied with the Principles of Laboratory Animal Care (NIH Publication No. 85–23, revised 1996) and were approved by the Institutional Ethics Committee (Approval No: IMSU/BCM/ETS/2024-09).

Animals were randomized into six groups (n = 4 per group, total 24 rats):

Group1: Control (normal chow, water only)

Group 2: Avocado only (avocado pulp supplementation)

Group 3: L-NAME (vascular injury model)

Group 4: L-NAME + drugs (metoprolol 25 mg/kg + losartan 20 mg/kg)

Group 5: L-NAME + avocado (80% w/w diet inclusion)

Group 6: L-NAME + drugs (metoprolol 25 mg/kg + losartan 20 mg/kg) + avocado (80% w/w inclusion).

### Induction of Vascular Injury and Interventions

Endothelial dysfunction was induced by oral administration of Nω-nitro-L-arginine methyl ester (L-NAME; 40 mg/kg/day in drinking water) for 4 weeks. Avocado pulp was freshly prepared daily, homogenized, and thoroughly incorporated into standard chow at 80% w/w to ensure uniform intake throughout the feeding period. Standard pharmacological therapy comprised oral metoprolol (25 mg/kg/day) and losartan (20 mg/kg/day). The treatment groups therefore received either avocado-supplemented diet, drug therapy, or their combination alongside L-NAME exposure, while controls received unsupplemented chow and water only. Morphometric Assessments Body weight, feed intake, and feed conversion efficiency (FCE) were measured weekly. Final body weight, body length, and body mass index (BMI; g/cm^2^) were recorded at sacrifice following 12h overnight fasting.

### Sample Collection

At study termination, animals were anesthetized with pentobarbital (40 mg/kg i.p.), and blood was collected by cardiac puncture. Serum was separated by centrifugation at 3,500 rpm for 15 min and stored at –20 °C until analysis.

### Biochemical Analyses

Serum lipid profile was quantified enzymatically using commercial kits: total cholesterol (TC), triglycerides (TG), LDL, HDL, and VLDL (Randox Laboratories, UK). Cardiac integrity was assessed via troponin I ELISA (Eastbiopharm, Hangzhou, China), creatine kinase (CK) activity by the IFCC-recommended method, and lactate dehydrogenase (LDH) activity using a colorimetric kit (Sigma-Aldrich, MAK066). Vascular biomarkers were measured using ELISA kits: endothelin-1, vascular cell adhesion molecule-1 (VCAM-1), and angiotensin II (AngII) (Elabscience, Wuhan, China). These methods are consistent with previously validated protocols in our laboratory.^15^

### Advanced Analytical Approaches

To interrogate systems-level interactions, we generated contour density plots of pairwise biomarker relationships and constructed correlation networks (nodes = biomarkers; edges = Spearman correlations with |ρ| ≥ 0.7, p < 0.05). Network topology metrics (density, clustering coefficient, average degree) were computed using NetworkX (Python).

### Statistical Analysis

Data are presented as mean ± SEM. One-way analysis of variance (ANOVA) followed by Tukey’s HSD test was used for group comparisons. Significance was set at p < 0.05. Graphical analyses were performed with GraphPad Prism v10 and Python (Matplotlib, Seaborn).

## 3.0: Results

### 3.1: Morphometric responses to avocado supplementation and pharmacological treatment in an L-NAME–induced vascular injury model

To evaluate the effects of avocado supplementation on overall growth performance, we compared morphometric indices across treatment groups (Figure 1a–f). Feed intake (FI) varied significantly among groups (ANOVA p = 0.0002), with the L-NAME group (Group 3) exhibiting the lowest consumption, consistent with impaired metabolic activity. In contrast, avocado-treated groups (Groups 2, 5, and 6) maintained intake levels comparable to controls, suggesting preserved appetite and dietary utilization. Weight gain (WG) was markedly suppressed in L-NAME-treated rats relative to controls and avocado-supplemented groups (p < 0.01), whereas the combination of avocado with standard drugs (Group 6) partially restored weight gain to near-control levels. Feed conversion efficiency (FCE) was profoundly reduced by L-NAME (Group 3), indicating impaired nutrient assimilation. Avocado supplementation, alone or in combination with drugs, significantly improved FCE (p < 0.01 versus Group 3), underscoring its role in maintaining metabolic efficiency. Final body weight followed a similar pattern (p < 0.0001), with L-NAME rats showing the greatest loss, while avocado supplementation mitigated weight decline in both single and combination regimens. Length measurements did not differ significantly among groups (p = 0.0566), reflecting that treatment effects were primarily metabolic rather than structural. However, body mass index (BMI) was significantly reduced in L-NAME animals (p = 0.0002), with avocado supplementation restoring BMI to levels comparable with control and drug-treated groups. Collectively, these results demonstrate that L-NAME–induced vascular injury is associated with reduced feed intake, diminished weight gain, and impaired nutrient utilization. Dietary avocado supplementation effectively counteracted these changes, improving FCE, weight, and BMI, with additive effects observed in combination with standard pharmacological therapy. These findings highlight the potential of avocado as a dietary adjunct for preserving morphometric and metabolic integrity under conditions of vascular stress.

**Figure 1.**
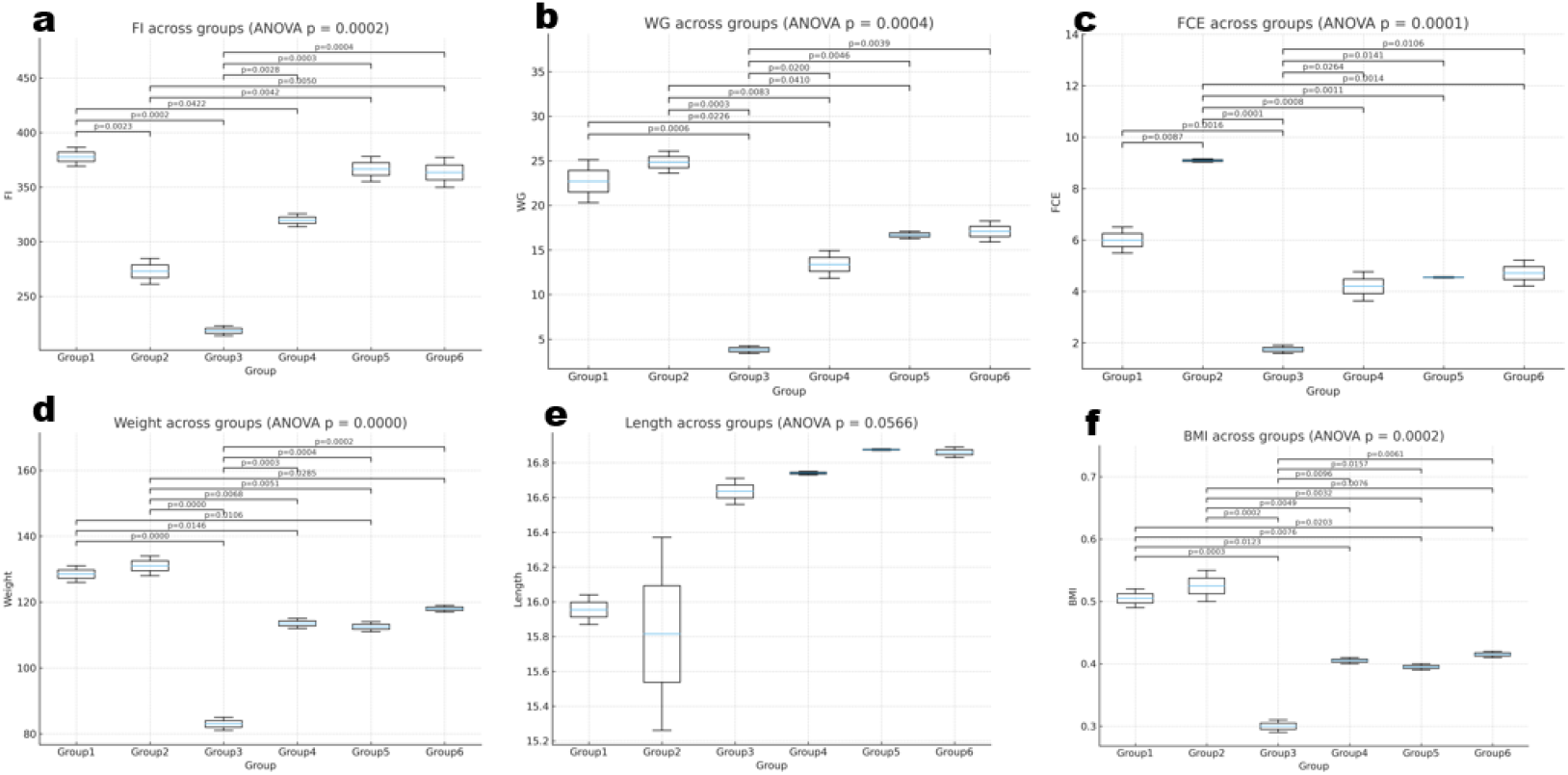
Effects of dietary avocado supplementation and pharmacological treatment on morphometric parameters in L-NAME–induced vascular injury model. Box plots show treatment group distributions (n = 4 per group) for (a) feed intake (FI), (b) weight gain (WG), (c) feed conversion efficiency (FCE), (d) final body weight, (e) body length, and (f) body mass index (BMI). Groups: Group 1 = Control (normal diet); Group 2 = Avocado only; Group 3 = L-NAME; Group 4 = L-NAME + drugs (metoprolol + losartan); Group 5 = L-NAME + avocado; Group 6 = L-NAME + drugs + avocado. Horizontal brackets indicate pairwise group comparisons with p-values derived from Tukey’s HSD post-hoc test following one-way ANOVA. Significant group differences (p < 0.05) are annotated.

**Figure 2.**
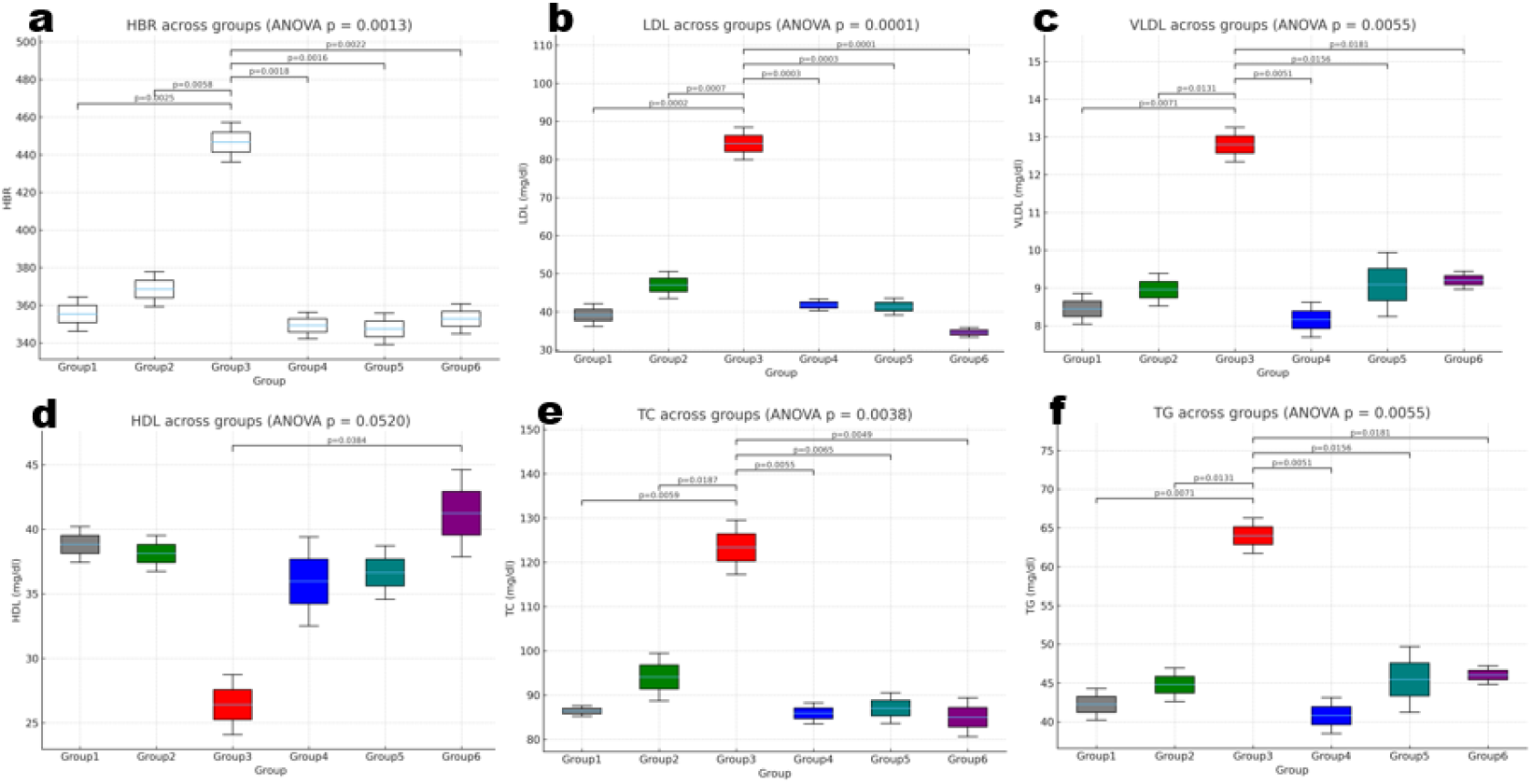
Effects of avocado supplementation and pharmacological treatment on cardiovascular and lipid profile parameters in L-NAME–induced vascular injury model. Box plots show treatment group distributions (n = 4 per group) for (a) heart beat rate (HBR), (b) low-density lipoprotein cholesterol (LDL), (c) very low-density lipoprotein cholesterol (VLDL), (d) high-density lipoprotein cholesterol (HDL), (e) total cholesterol (TC), and (f) triglycerides (TG). Groups: Group 1 = Control (normal diet); Group 2 = Avocado only; Group 3 = L-NAME; Group 4 = L-NAME + drugs (metoprolol + losartan); Group 5 = L-NAME + avocado; Group 6 = L-NAME + drugs + avocado. Horizontal brackets indicate pairwise group comparisons with p-values derived from Tukey’s HSD post-hoc test following one-way ANOVA. Significant differences (p < 0.05) are annotated.

**Figure 3.**
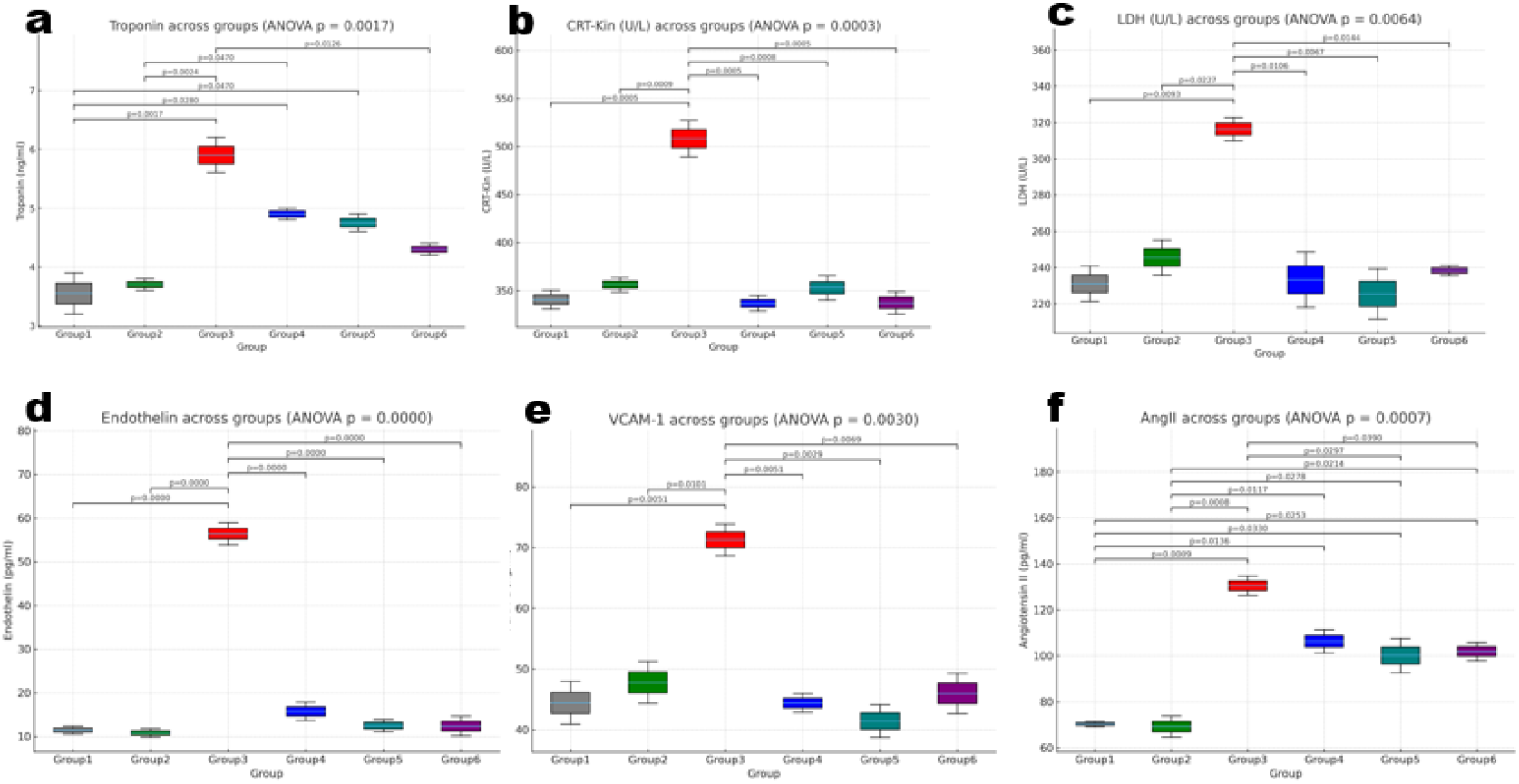
Effects of avocado supplementation and pharmacological treatment on cardiac integrity and vascular injury markers in L-NAME–induced vascular damage model. Box plots show treatment group distributions (n = 4 per group) for (a) troponin, (b) creatine kinase (CRT-Kin), (c) lactate dehydrogenase (LDH), (d) endothelin, (e) vascular cell adhesion molecule-1 (VCAM-1), and (f) angiotensin II (AngII). Groups: Group 1 = Control (normal diet); Group 2 = Avocado only; Group 3 = L-NAME; Group 4 = L-NAME + drugs (metoprolol + losartan); Group 5 = L-NAME + avocado; Group 6 = L-NAME + drugs + avocado. Horizontal brackets indicate pairwise group comparisons with p-values derived from Tukey’s HSD post-hoc test following one-way ANOVA. Significant group differences (p < 0.05) are annotated.

### 3.2: Cardiovascular and lipid profile responses to avocado supplementation and pharmacological treatment in an L-NAME-induced vascular injury model

Heartbeat rate (HBR) was significantly altered across groups (ANOVA p = 0.0013). Rats treated with L-NAME (Group 3) exhibited the highest HBR, consistent with hypertension-induced tachycardia. Avocado supplementation, either alone (Group 2) or in combination with pharmacological therapy (Groups 5 and 6), normalized HBR to near-control levels, demonstrating a cardioprotective effect. Lipid profiles revealed marked dyslipidemia in the L-NAME group. LDL cholesterol was significantly elevated in Group 3 compared with controls (p < 0.001), while both avocado-supplemented and drug-treated groups displayed reduced LDL-c concentrations, indicating partial restoration of lipid homeostasis. Similarly, VLDL-c and triglyceride (TG) levels were highest in the L-NAME group (p < 0.01), with significant reductions observed in avocado-supplemented and drug-treated animals. Total cholesterol (TC) followed the same pattern, with L-NAME markedly elevating TC (p = 0.0038), whereas avocado and combination therapy normalized the values. HDL cholesterol, by contrast, was reduced in the L-NAME group, reflecting impaired reverse cholesterol transport. Although the ANOVA result approached significance (p = 0.0520), avocado supplementation was associated with modest HDL preservation, particularly in Group 6 (L-NAME + drugs + avocado), which showed the highest HDL levels among the treated groups. Overall, these findings indicate that L-NAME induces pronounced dyslipidemia characterized by elevated LDL, VLDL, TC, TG, and reduced HDL. Avocado supplementation effectively countered these disturbances, restoring lipid balance either alone or synergistically with pharmacological therapy. The normalization of both HBR and lipid parameters underscores the dual role of avocado in cardiovascular protection, combining hemodynamic stabilization with correction of lipid abnormalities.

### 3.3: Effects of avocado supplementation on cardiac integrity and vascular injury markers in an L-NAME–induced vascular injury model

Markers of cardiac damage were strongly elevated in L-NAME–treated rats. Troponin, a sensitive indicator of myocardial injury, was significantly higher in Group 3 compared with controls (p = 0.0017). Avocado supplementation (Groups 2 and 5) and drug treatment (Group 4) reduced troponin levels, with the combination therapy group (Group 6) showing values approaching baseline. Similarly, creatine kinase (CRT-Kin) and lactate dehydrogenase (LDH) were elevated by L-NAME (p = 0.0003 and p = 0.0064, respectively), reflecting cardiomyocyte membrane leakage. Both avocado and drug treatments attenuated these increases, with the most pronounced reductions observed in the avocado + drug group. Vascular injury markers followed a comparable trend. Endothelin, a potent vasoconstrictor, was markedly elevated in L-NAME animals (p < 0.0001), confirming endothelial dysfunction. Avocado supplementation significantly reduced endothelin concentrations, indicating improved vascular tone. VCAM-1, a key adhesion molecule upregulated during vascular inflammation, was also elevated in L-NAME (p = 0.0030). Avocado treatment reduced VCAM-1 levels toward control values, suggesting mitigation of endothelial activation. Angiotensin II (AngII), central to the renin–angiotensin system, was highest in L-NAME–treated rats (p = 0.0007). Avocado supplementation, particularly in Groups 5 and 6, lowered AngII concentrations, indicating a dampening of vasoconstrictive and pro-inflammatory signaling. Taken together, these results demonstrate that avocado supplementation provides robust cardioprotective and vasculoprotective effects. By reducing biomarkers of myocardial injury (troponin, CK, LDH) and attenuating vascular dysfunction (endothelin, VCAM-1, AngII), avocado not only preserved cardiac integrity but also alleviated endothelial stress, reinforcing its potential as a dietary adjunct in the management of vascular injury.

### 3.4: Pairwise contour analysis of integrated biomarker relationships

To further interrogate how treatment regimens altered biomarker interactions, we applied contour-based visualization of pooled kernel density estimates (KDE) across groups (Figure 4a–d). LDL and TG exhibited a strong positive association, with the highest clustering observed in L-NAME–treated rat (Group 3), reflecting a dyslipidemic phenotype characterized by simultaneous hypercholesterolemia and hypertriglyceridemia. Avocado-supplemented groups (Groups 2, 5, and 6) shifted toward lower-density zones comparable to controls, indicating lipid normalization. Heart beat rate (HBR) versus endothelin demonstrated that L-NAME rats occupied the extreme upper quadrant, consistent with simultaneous tachycardia and endothelial dysfunction (Figure 4b). In contrast, avocado supplementation reduced both HBR and endothelin levels, returning animals to contour regions overlapping with control values. Cardiac stress biomarkers revealed a strong co-elevation of troponin and angiotensin II in L-NAME animals (Figure 4c). This clustering was unique to the pathological state, with avocado or drug treatments attenuating the distribution toward lower-density zones. The restoration of troponin–AngII coupling to near-control levels highlights the cardioprotective effects of avocado. Finally, LDL and HDL exhibited an inverse relationship (Figure 4d). L-NAME rats clustered at high LDL and low HDL values, consistent with impaired reverse cholesterol transport. Avocado-containing groups showed a shift toward higher HDL and reduced LDL, again approximating control distributions. Together, these contour plots underscore the integrative nature of L-NAME–induced pathology, wherein dyslipidemia, endothelial dysfunction, and cardiac stress converge, and demonstrate the ability of avocado supplementation to remodel these pathological biomarker interactions toward healthier phenotypic patterns.

**Figure 4.**
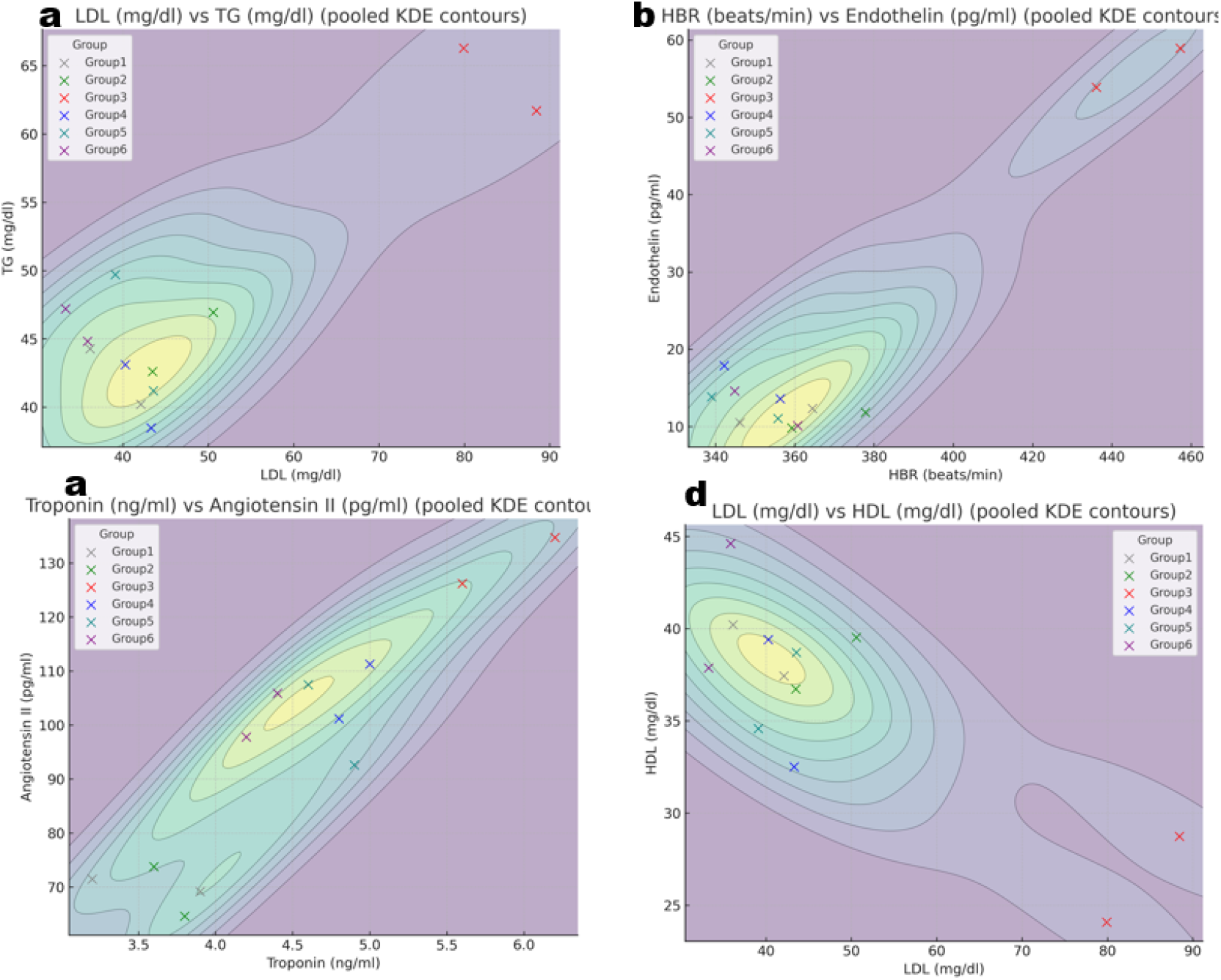
Contour density plots of pairwise biomarker relationships across treatment groups. Kernel density estimation (KDE) contour plots illustrate pooled distributions of selected biomarker pairs with individual group replicates overlaid (n = 4 per group). Panels show (a) LDL vs TG, (b) heart beat rate (HBR) vs endothelin, (c) troponin vs angiotensin II (AngII), and (d) LDL vs HDL. Contour shading represents increasing density of observations, while colored markers indicate group assignments: Group 1 = Control (gray); Group 2 = Avocad only (green); Group 3 = L-NAME (red); Group 4 = L-NAME + drugs (blue); Group 5 = L-NAME + avocado (teal); Group 6 = L-NAME + drugs + avocado (purple). These plots highlight how L-NAME treatment drives clustering in pathological biomarker zones (e.g., high LDL and TG, high HBR with endothelin, elevated troponin with AngII, reduced HDL), whereas avocado supplementation and drug combinations shift distributions toward control-like regions.

### 3.5: VCAM-1 correlations with cardiac injury and lipid markers

We next examined vascular inflammation by analyzing VCAM-1 relationships with cardiac enzymes and lipid parameters (Figure 5a–d). VCAM-1 exhibited strong positive coupling with LDH (Figure 5a) and CK (Figure 5b), particularly in L-NAME–treated animals, consistent with endothelial activation paralleling cardiomyocyte injury. These associations were attenuated in avocado-supplemented groups, which clustered at lower LDH and CK levels, indicating reduced cardiac leakage. Similarly, VCAM-1 showed direct associations with triglycerides (Figure 5c) and total cholesterol (Figure 5d). L-NAME rats were uniquely clustered in zones of high VCAM-1 with elevated TG and TC, reflecting intertwined dyslipidemia and vascular inflammation. By contrast, avocado supplementation, alone or with drugs, shifted distributions toward lower VCAM-1 and lipid values, resembling control patterns. These findings highlight VCAM-1 as a central integrator linking vascular inflammation with both lipid dysregulation and cardiac injury in L-NAME pathology. Avocado supplementation effectively decoupled these pathological relationships, reducing endothelial activation and restoring biomarker interactions toward physiologic norms.

**Figure 5.**
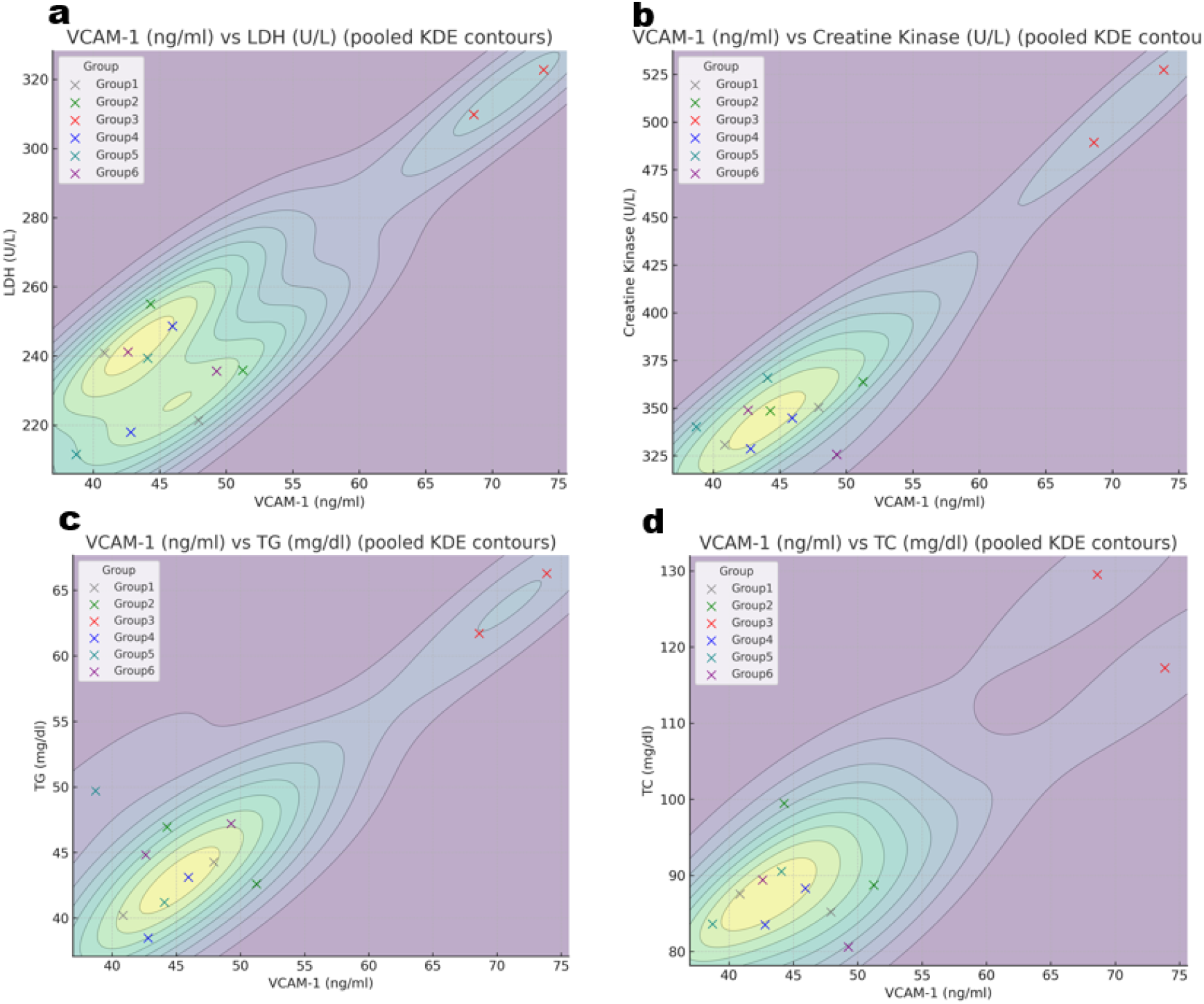
Contour density plots of VCAM-1 in relation to cardiac integrity and lipid markers across treatment groups. Kernel density estimation (KDE) contour plots show pooled distributions of VCAM-1 against selected cardiac and lipid biomarkers, with individual group replicates overlaid (n = 4 per group). Panels display (a) VCAM-1 vs LDH, (b) VCAM-1 vs creatine kinase (CK), (c) VCAM-1 vs triglycerides (TG), and (d) VCAM-1 vs total cholesterol (TC). Contour shading represents increasing density of observations, while colored points denote treatment groups: Group 1 = Control (gray); Group 2 = Avocado only (green); Group 3 = L-NAME (red); Group 4 = L-NAME + drugs (blue); Group 5 = L-NAME + avocado (teal); Group 6 = L-NAME + drugs + avocado (purple). These plots highlight strong pathological clustering of L-NAME–treated rats (red) at high VCAM-1 levels alongside elevated LDH, CK, TG, and TC, consistent with vascular inflammation, cardiac damage, and dyslipidemia. Avocado supplementation, alone or in combination with drugs, shifted these distributions toward control-like patterns, reflecting attenuation of endothelial activation and improved cardiometabolic balance.

## 4.0: Discussion

This study shows that dietary avocado supplementation exerts broad protective effects in the setting of L-NAME-induced vascular injury, restoring metabolic balance, lipid homeostasis, cardiac integrity, and vascular function. Across morphometric, biochemical, and network-level analyses, avocado consistently mitigated pathological changes and shifted biomarker interactions toward control-like patterns, underscoring its potential as a systems-level dietary intervention. Morphometric outcomes provided early evidence of benefit. L-NAME reduced feed intake, weight gain, feed conversion efficiency, and BMI, indicating impaired nutrient assimilation and metabolic stress. Avocado supplementation reversed these deficits, restoring BMI and weight gain without affecting body length. This suggests improved nutrient utilization rather than structural growth, likely mediated by avocado’s monounsaturated fatty acids, phytosterols, and antioxidant bioactives that preserve metabolic efficiency under stress.^11,16^ Lipid and cardiovascular markers further demonstrated avocado’s protective capacity. L-NAME induced dyslipidemia with elevated LDLc, VLDLc, total cholesterol, and triglycerides, coupled with reduced HDLc, a profile strongly associated with vascular injury.^17^ Avocado countered these effects, lowering atherogenic lipoproteins and partially preserving HDL, while also normalizing the L-NAME-induced increase in heart rate. Together, these findings highlight avocado’s dual role in correcting lipid abnormalities and alleviating hemodynamic stress.^18^ Markers of cardiac integrity confirmed myocardial protection. L-NAME elevated troponin, creatine kinase, and lactate dehydrogenase, consistent with cardiomyocyte injury, and increased angiotensin II, reflecting maladaptive neurohormonal activation.^19^ Avocado supplementation attenuated these rises, particularly when combined with pharmacological therapy, restoring cardiac stress markers toward baseline. The reduction of angiotensin II is especially notable, as it suggests modulation of renin-angiotensin signaling, a critical driver of vascular and myocardial remodeling.^20^ Vascular inflammation emerged as a central feature of L-NAME pathology. Endothelin and VCAM-1 were markedly elevated, and VCAM-1 showed strong correlations with lipid and cardiac injury markers, linking dyslipidemia and myocardial damage to endothelial dysfunction.^21,22^ Avocado lowered VCAM-1 and disrupted these pathological associations, effectively decoupling vascular inflammation from metabolic and cardiac injury. Beyond individual biomarkers, contour and network analyses revealed avocado’s impact on systems-level organization. L-NAME produced pathological clustering of biomarkers, high LDLc coupled to triglycerides, elevated heart rate with endothelin, troponin tightly aligned with angiotensin II, and reflecting maladaptive integration. Avocado supplementation shifted distributions toward control-like densities, disrupted pathological couplings, and simplified network architecture. Network metrics confirmed that L-NAME increased correlation density and clustering, while avocado restored simpler, physiologic networks. These effects indicate that avocado not only alters biomarker levels but also reprograms the connectivity of biological systems. Together, these findings position avocado as a nutritionally rich food capable of countering endothelial dysfunction, dyslipidemia, and cardiac stress through integrated mechanisms. By improving metabolic efficiency, restoring lipid balance, protecting cardiomyocytes, and attenuating vascular inflammation, avocado simplified pathological biomarker networks and re-established control-like physiological coherence.^22,23^ Such systems-level remodeling highlights avocado’s potential as a nutraceutical adjunct in strategies for preventing or mitigating cardiovascular disease.

## Conclusion

In summary, avocado supplementation attenuated the systemic consequences of L-NAME–induced vascular injury, improving morphometric efficiency, normalizing lipid profiles, reducing cardiac injury markers, and alleviating endothelial dysfunction. Beyond correcting individual biomarkers, avocado disrupted maladaptive couplings, simplified pathological correlation networks, and restored control-like patterns of physiological integration. These findings highlight avocado as more than a nutritional supplement: it functions as a systems-level modulator that stabilizes cardiovascular and metabolic homeostasis under stress. The breadth of effects observed suggests translational potential for avocado as a nutraceutical adjunct in preventing or managing cardiometabolic diseases characterized by endothelial activation, dyslipidemia, and myocardial stress. Future work in larger cohorts and clinical settings will be essential to validate these findings and define mechanisms, but the present results provide compelling preclinical evidence for the integrative cardioprotective benefits of avocado.

## Limitations and Future directions

While the present findings establish avocado pulp as a potent dietary modulator of vascular and cardiac injury, several questions warrant further exploration. Larger preclinical cohorts are needed to validate these effects with greater statistical power and to delineate dose-response relationships. Mechanistic studies should focus on identifying the bioactive compounds responsible for avocado’s cardioprotective actions, particularly their influence on nitric oxide signaling, lipid metabolism, and inflammatory pathways. Integrative omics approaches-including transcriptomics, metabolomics, and lipidomics could reveal how avocado reprograms molecular networks underlying endothelial function and myocardial stress. Clinical translation will require controlled dietary intervention trials in humans at risk for hypertension and dyslipidemia, testing whether avocado-enriched diets confer similar systems-level protection. Finally, comparative studies with other functional foods may define avocado’s unique contributions within cardioprotective dietary patterns. Together, these directions will clarify avocado’s potential as a nutraceutical adjunct in cardiovascular prevention and therapy.

## Acknowledgements

We are grateful to the entire students of Departments of Biochemistry, and Human Nutrition, for their immense support. We appreciate the effort of Professor E.N. Agomuo as well as the IMSU animal facility for providing logistical support.

